# Novel phenotype of Wolbachia strain wPip in Aedes aegypti challenges assumptions on mechanisms of Wolbachia-mediated dengue virus inhibition

**DOI:** 10.1101/2020.02.20.957423

**Authors:** Johanna E. Fraser, Tanya B. O’Donnell, Johanna M. Duyvestyn, Scott L. O’Neill, Cameron P. Simmons, Heather A. Flores

## Abstract

The bacterial endosymbiont *Wolbachia* is a biocontrol tool that inhibits the ability of the *Aedes aegypti* mosquito to transmit positive-sense RNA viruses such as dengue and Zika. Growing evidence indicates that when *Wolbachia* strains *w*Mel or *w*AlbB are introduced into local mosquito populations, human dengue incidence is reduced. Despite the success of this novel intervention, we still do not fully understand how *Wolbachia* protects mosquitoes from viral infection. Here, we demonstrate that the *Wolbachia* strain *w*Pip does not inhibit virus infection in *Ae. aegypti*. We have leveraged this novel finding, and a panel of *Ae. aegypti* lines carrying virus-inhibitory (*w*Mel and *w*AlbB) and non-inhibitory (*w*Pip) strains in a common genetic background, to rigorously test a number of hypotheses about the mechanism of *Wolbachia*-mediated virus inhibition. We demonstrate that, contrary to previous suggestions, there is no association between a strain’s ability to inhibit dengue infection in the mosquito and either its typical density in the midgut or salivary glands, or the degree to which it elevates innate immune response pathways in the mosquito. These findings, and the experimental platform provided by this panel of genetically comparable mosquito lines, clear the way for future investigations to define how *Wolbachia* prevents *Ae. aegypti* from transmitting viruses.

**Author summary:** Dengue virus, transmitted by the *Aedes aegypti* mosquito, is one of the fastest-growing infectious diseases, causing an estimated 390 million human infections per year worldwide. Vaccines have limited efficacy and there are no approved therapeutics. This has driven the rise of novel vector control programs, in particular those that use the bacterium, *Wolbachia*, which prevents transmission of dengue and other human pathogenic viruses when stably introduced into *Ae. aegypti* populations. Although this is proving to be a highly effective method, the details of how this biocontrol tool works are not well understood. Here we characterise a new *Wolbachia* strain, *w*Pip, and find that *Ae. aegypti* carrying *w*Pip are still able to transmit dengue similar to mosquitoes that do not carry *Wolbachia*. This finding has allowed us to begin a rigorous program of comparative studies to determine which features of a *Wolbachia* strain determine whether it is antiviral. Understanding these mechanisms will enable us to predict the risk of viral resistance arising against *Wolbachia* and facilitate preparation of second-generation field release lines.

## Introduction

The *Aedes aegypti* mosquito is the primary vector for many human pathogenic viruses including dengue (DENV), Zika (ZIKV) and chikungunya (CHIKV). Global incidence of arthropod-borne viruses (arboviruses) such as these is increasing in response to urbanization in the tropics, as well as globalization and expansion of the geographical range of *Ae. aegypti* [1, 2]. Arboviruses like DENV are typically associated with acute, self-limiting febrile disease, although severe manifestations can occur, in some instances leading to death. There are currently no approved specific antiviral therapeutics for DENV, ZIKV or CHIKV and recently developed vaccines for DENV are suboptimal and controversial [3-6]. As such, treatment is supportive only and limiting virus transmission is largely dependent on vector control. Conventional vector control programs attempt to suppress mosquito populations by removing breeding sites, or using insecticide sprays targeting adult mosquito habitats. However, the increasing incidence of these diseases demonstrates that current control programs are failing and there is a need for novel efficacious and cost-effective alternatives.

A new generation of vector control approaches exploits the obligate endosymbiont, *Wolbachia pipientis*. Various strains of *Wolbachia* are naturally found in at least 40% of insect species [7]. *Wolbachia* is maternally transmitted, and some strains induce a phenomenon known as cytoplasmic incompatibility (CI) which provides a reproductive advantage to females that carry *Wolbachia*. Significantly, many *Wolbachia* strains can protect arthropod hosts from viral infection [8-11]. *Wolbachia* is not typically found in *Ae. aegypti*, but it can be stably transferred into these mosquitoes by microinjection. Some *Wolbachia* strains including *w*Mel (derived from *Drosophila melanogaster*) and *w*AlbB (from *Ae. albopictus*), protect *Ae. aegypti* from infection by flaviviruses such as DENV and ZIKV, and alphaviruses including CHIKV [11-14]. Recent field trials have tested the utility of these mosquitoes as a biocontrol method. Aided by maternal transmission, as well as CI, short term releases of *w*Mel- or *w*AlbB-carrying *Ae. aegypti* have resulted in rapid introgression into wild *Ae. aegypti* populations. Recent reports indicate this can significantly reduce DENV incidence [15-17].

Despite success of these field trials, we do not understand how *Wolbachia* inhibits arbovirus infection. Several hypotheses have been proposed, including competition between *Wolbachia* and viruses for metabolic resources and physical space within host cells [11, 18-20], modification of host gene expression, and immune priming in new hosts [11, 21-23]. Understanding the contributions of each of these mechanisms to arboviral inhibition in *Wolbachia*-carrying *Ae. aegypti* has proven to be complex, and it is likely that the antiviral syndrome that *Wolbachia* species induce in the host is multi-layered, with no single mechanism responsible for this phenotype [24, 25].

We and others have produced and characterized a series of *Ae. aegypti* lines transinfected with different *Wolbachia* strains (summarized in Table 1), including several strains from *Drosophila* species, and different mosquito species.

**Table 1.** *Wolbachia* stains that have been introduced into *Ae. aegypti* to date.

| Host species | *Wolbachia* strain(s) | *Wolbachia* supergroup | References |
| --- | --- | --- | --- |
| *D. melanogaster* | wMel, wMelCs, wMelPop | A | [26-28] |
| *D. simulans* | wRi, wAu | A | [26, 29] |
| *Ae. albopictus* | wAlbA, wAlbB | A and B, respectively | [29-31] |
| *Cx. quinquefasciatus* | wPip | B | [26] |

So far, all 7 of the *Wolbachia*-carrying *Ae. aegypti* lines tested have provided some level of protection against flaviviruses. Here, we characterize the vector competence of *Ae. aegypti* transinfected with *w*Pip (from *Culex quinquefasciatus*, supergroup B). Past data has suggested that removal of *w*Pip from its natural host by antibiotic treatment leads to an increase in the replication of DENV-related flavivirus West Nile virus (WNV), indicating that *w*Pip may be antiviral in this context [32]. We previously reported the generation of a *w*Pip-*Ae. aegypti* line [26], and that *w*Pip resides at a high density in *Ae. aegypti* – a feature widely regarded as important for *Wolbachia* to impart its antiviral effects [33-35]. However, we show here that *w*Pip does not restrict flavivirus replication, dissemination or reduce the transmission potential in *Ae. aegypti*. We leverage this finding to understand more about what makes a *Wolbachia* strain antiviral. We compare the density and tissue specificity of *w*Pip and antiviral strains *w*Mel and *w*AlbB, in *Ae. aegypti* backcrossed onto the common laboratory Rockefeller *Ae. aegypti* line and identify no link between strains that reside at high density in the midgut or salivary glands, and those that protect against DENV. Furthermore, we determine that commonly used proxies of immune activation of the host are not induced by all antiviral *Wolbachia* strains. Thus, our finding of a *Wolbachia* strain that does not inhibit DENV replication in *Ae. aegypti* has facilitated the first rigorous tests to determine the mechanisms that drive *Wolbachia*-mediated virus inhibition in this species.

## Results

### *w*Pip does not block flavivirus replication in virus-injected *Ae. aegypti*

Past reports suggest *w*Pip is a good antiviral candidate for testing in *Ae. aegypti*, as it may provide viral protection to *Cx. quinquefasciatus* [32], and it resides at high density in *Ae. aegypti* [26]. We assessed the vector competence of *w*Pip-*Ae. aegypti* by intrathoracic injection challenge and infectious blood meal, comparing DENV replication in this line to a matched *Wolbachia*-free line (tetracycline-treated *w*Pip-*Ae. aegypti*; *w*Pip.Tet). As a control, we included *w*Mel-*Ae. aegypti* (predominantly used by the World Mosquito Program as their field release line), and its matched *Wolbachia*-free line (*w*Mel.Tet).

Briefly, 60 seven-day old females were injected with 6.3 × 10^5^ TCID_50_/ml of DENV-3, or a 10-fold dilution thereof. Total RNA was extracted from whole, surviving mosquitoes 7 days post infection. Absolute DENV-3 RNA copies were determined in each mosquito by qRT-PCR and extrapolation to a standard curve. Consistent with previous reports, *w*Mel restricted DENV replication by approximately 1log_10_ compared to its matched Tet control line, when injected with either concentration of virus (Fig. 1A and B) [26, 36]. Notably, 37 of 49 injected *w*Mel mosquitoes scored positive for DENV-3 infection (>1000 copies/mosquito) when injected with 6.3 × 10^5^ TCID_50_/ml, compared to just 15 of 47 *w*Mel mosquitoes when injected with 6.3 × 10^4^ TCID_50_/ml (Fig. 1A and B, in parenthesis above the bar charts), consistent with better viral restriction occurring when the mosquitoes are challenged with a lower virus titre [26]. In stark contrast to the protection afforded by *w*Mel, *w*Pip did not inhibit either concentration of DENV-3 compared to the matched *w*Pip.Tet control line. As well as failing to reduce the amount of virus in these mosquitoes, the number of infected *w*Pip-mosquitoes also remained high and comparable to the matched *w*Pip.Tet control line, demonstrating this *Wolbachia* strain does not provide antiviral protection when transinfected into *Ae. aegypti*.

**Fig 1.**
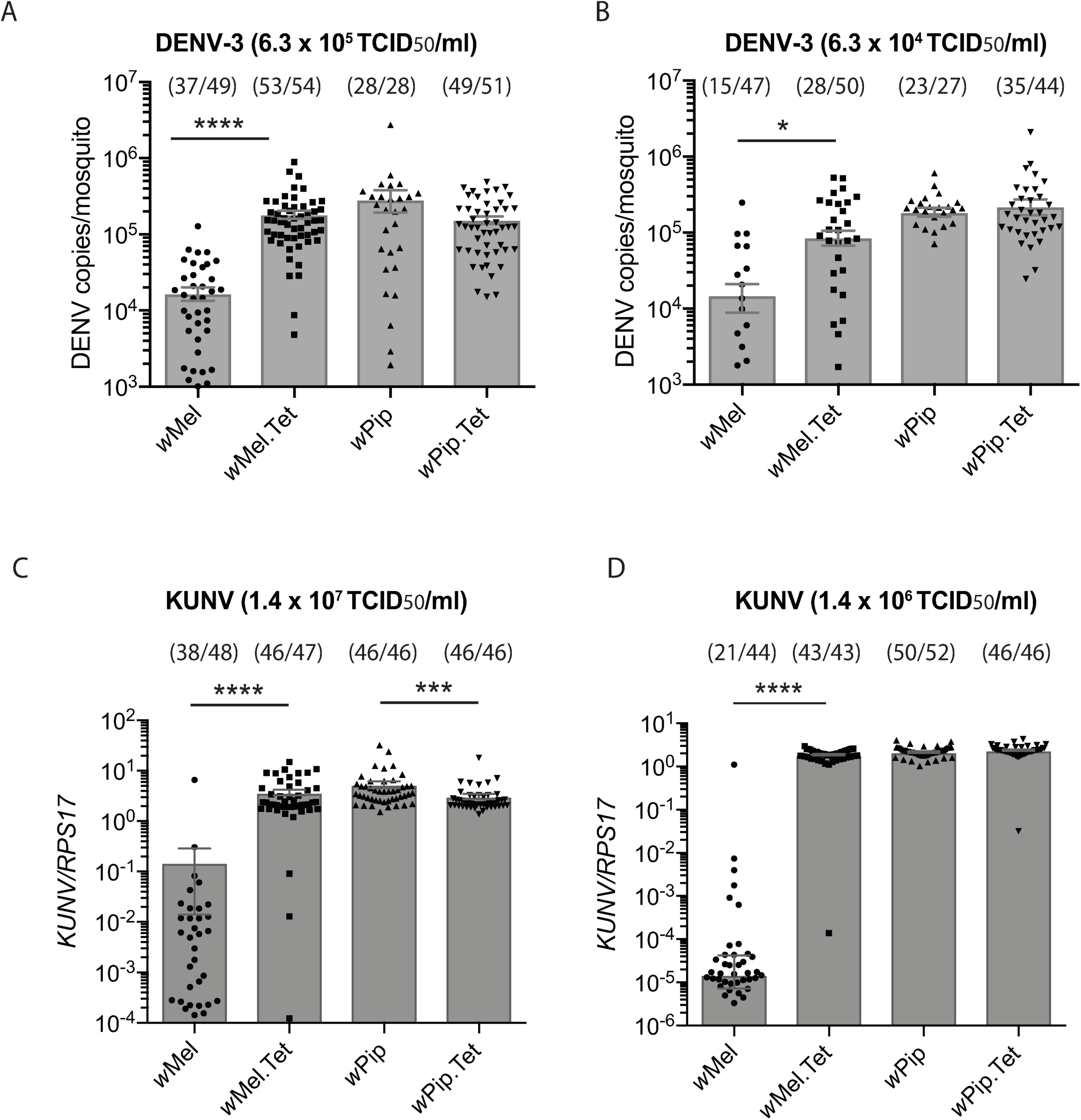
*w*Pip does not inhibit flavivirus replication in virus-injected *Ae. aegypti*. Intrathoracic injections of 6 or 7-day old female mosquitoes were performed: (A) DENV-3 at 6.3 ×10^5^ TCID_50_/ml or (B) 6.3 ×10^4^ TCID_50_/ml, and (C) KUNV at 1.4 × 10^7^ TCID_50_/ml or (D) 1.4 × 10^6^ TCID_50_/ml. RNA was extracted from whole mosquito bodies 7-days post infection and virus replication was quantified by qRT-PCR. Data are the mean number of virus genome copies per mosquito (DENV) or per *rps17* mosquito house-keeping gene (KUNV) ± SEM with individual data points overlaid. Data are representative of 2 independent experiments. Number of DENV-3/KUNV positive mosquitoes/total injected mosquitoes are indicated above each bar. Statistical analyses were performed using a Mann-Whitney test where * p < 0.05, ***p<0.001, ****p<0.0001.

Since *w*Pip may inhibit WNV in its native host mosquito species, *Cx. quinquefasciatus* [32], we next examined whether *w*Pip could inhibit WNV in an *Ae. aegypti* host. *Ae. aegypti* carrying *w*Pip, *w*Mel, or their respective Tet control lines, were injected with the non-pathogenic WNV strain Kunjin virus (KUNV) at 1.4 x 10^7^ TCID_50_/ml or a 10-fold dilution thereof. Mosquitoes were collected 7-days post-infection and RNA was isolated from whole mosquitoes. Total KUNV RNA copies relative to mosquito host *RPS17* RNA, was determined in each mosquito by qRT-PCR. Similar to previous reports, *w*Mel provided substantial protection against KUN, with a nearly 2log_10_ reduction in viral RNA copies measured in *w*Mel-infected mosquitoes compared to *w*Mel.Tet (Fig. 1 C and D) [36]. However, as was observed for DENV-3 infections, *w*Pip failed to provide protection against KUNV at either high or low concentrations of virus. Thus, *w*Pip is not antiviral towards flaviviruses in *Ae. aegypti*.

### *w*Pip does not restrict DENV replication, dissemination or transmission in *Ae. aegypti* following an infectious blood meal

Since the route of viral infection can affect the ability of a *Wolbachia* strain to inhibit flaviviruses [30], we next examined the impact of *w*Pip on *Ae. aegypti* infection by DENV following an infectious blood meal. Female mosquitoes carrying *w*Mel or *w*Pip and their respective Tet control lines were fed a blood meal containing freshly harvested cell culture-derived DENV-3 (6.6 × 10^6^ TCID_50_/mL diluted 1:1 in sheep blood). Mosquitoes were incubated for 15 days, then the body and head of the mosquito were collected separately as indicators of established infection and viral dissemination, respectively. No *w*Mel-carrying mosquitoes that took a blood meal established a DENV-3 infection, compared to 74% of the matched Tet control cohort, and a >4log_10_ reduction in mean DENV-3 copies/body were observed in the presence of *w*Mel (Fig. 2 and Table 2). By contrast, 93% of *w*Pip-transinfected mosquito bodies scored positive for DENV-3, similar to 96% of the matched Tet control cohort, with a slight, although significant, decrease in the mean viral copies/mosquito in *w*Pip-carrying mosquitoes compared to its matched Tet control cohort. No *w*Mel mosquitoes had viral RNA disseminated to their heads, compared to 75% of *w*Mel.Tet mosquitoes. Similar numbers of *w*Pip- and *w*Pip.Tet-carrying mosquitoes scored positive for disseminated infection (83% and 97%, respectively), and the mean DENV copies/head was slightly but significantly greater in *w*Pip-infected mosquitoes compared to *w*Pip.Tet. Note that the slight increases and decreases in viral copy number observed for *w*Pip-carrying mosquitoes compared to the Tet control line are likely to be due to biological variability, as they were not consistent across 3 independent experiments. The overall trend clearly showed no difference between the vector competence of these two lines.

**Table 2.** Restriction of DENV-3 infection and dissemination by wPip.

|  | Body | Head | Saliva |
| --- | --- | --- | --- |
| Line | % DENV +<sup>#</sup><br>(*n*= 72) | % DENV +<sup>#</sup><br>(*n*= 72) | Donor mosquitoes with<br>infectious saliva (*n*) |
| wMel | 0 | 0 | 0 (15) |
| wMel.Tet | 74 | 75 | 6 (14) |
| wPip | 93 | 83 | 3 (14) |
| wPip.Tet | 96 | 97 | 5 (13) |

<sup>#</sup> Calculated as a percentage of total engorged mosquitoes.
(*n*) total mosquitoes examined.

**Fig 2.**
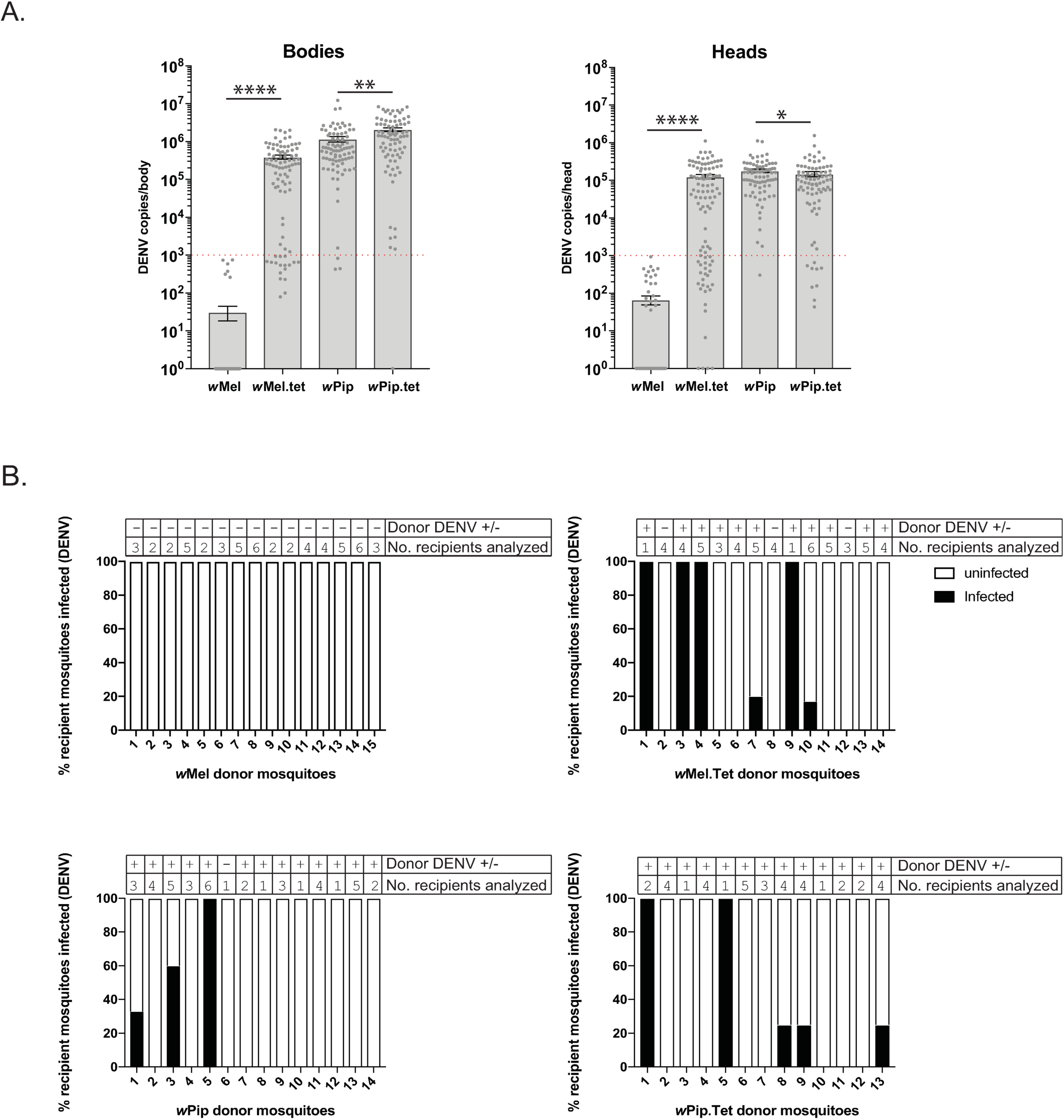
*w*Pip does not restrict DENV replication, dissemination or transmission in *Ae. aegypti* following an infectious blood meal. (A) Seven-day old female mosquitoes were fed a blood meal containing DENV-3 (6.6 × 10^6^ TCID_50_/ml) and incubated for 15 days. DENV genome copies were determined by qRT-PCR for each body as a measure of infection, and for each head as a measure of viral dissemination. Data are the mean viral genome copies per mosquito body or head ± SEM, with individual data points overlaid, and are representative of 3 independent experiments. Statistical analyses were performed using a Mann-Whitney test where * p < 0.05, **p < 0.01, ****p < 0.0001. Red line indicates LOD_95_ of the qRT-PCR reaction. Zero values have been plotted as 10^0^ (1) to allow visualization on the log_10_ scale (B). At 14 d.p.i., saliva from 15 mosquitoes/line from (A) was collected (donor mosquitoes) and each saliva sample injected into 6 *w*Mel.Tet mosquitoes (recipients) to determine whether the blood-fed mosquitoes were producing infectious virus. DENV genome copies in entire recipient mosquitoes were determined as in (A) and mosquitoes with DENV values above LOD_95_ were scored positive. Columns represent mosquitoes from a single donor, where black indicates the % recipients infected from a single donor mosquito. White is uninfected. Number of recipients analyzed from each donor, and whether the donor body was positive (+) or negative (-) for DENV are indicated above the columns.

We further examined the saliva from a proportion of mosquitoes that were fed a virus-spiked blood meal to determine whether infectious virus could be transmitted by *w*Pip-carrying mosquitoes. Saliva was collected from ∼15 mosquitoes/line (donor mosquitoes) at 15-days post-infection. Each saliva sample was then injected into the thorax of 6 *w*Mel.Tet recipient mosquitoes to assess the replication competence of the virus. Seven days later, injected mosquitoes were harvested, total RNA was extracted and qRT-PCR performed to determine the number of recipients positive for virus infection (>10^3^DENV copies/mosquito). *w*Mel caused a significant reduction in the number of donor mosquitoes that carried infectious DENV in their saliva, compared to its matched Tet control line (Fisher’s exact test p<0.05), but *w*Pip did not (Fig. 2B and Table 2).

These data consistently demonstrate that *w*Pip does not provide *Ae. aegypti* mosquitoes with protection against DENV-3 infection, dissemination or transmission.

### Differences in mosquito host genetics do not underlie *w*Pip’s lack of antiviral activity

Following microinjection of a new *Wolbachia* strain into *Ae. aegypti*, an intense genetic bottlenecking occurs as we select individual mosquitoes that carry *Wolbachia*. To determine whether the lack of antiviral activity observed for *w*Pip was due to specific genetic features of the host mosquito selected during this process, we backcrossed our *Wolbachia*-carrying lines to the inbred laboratory mosquito line, Rockefeller, through six generations, placing each *Wolbachia* strain into the same nuclear genetic background. Intrathoracic injection of these lines with DENV-2 confirmed a lack of viral restriction by *w*Pip (Supp Fig. 1). Thus, the inability of *w*Pip-*Ae. aegypti* to inhibit flaviviruses is consistent in different *Ae. aegypti* nuclear backgrounds.

### High *Wolbachia* density in *Ae. aegypti* salivary glands and midgut is not required for viral inhibition

Our novel identification of a *Wolbachia* strain that does not appear to restrict flaviviruses in *Ae. aegypti*, generated an opportunity to determine what features of *Wolbachia* are common to antiviral strains. Several reports have suggested that the ability of a *Wolbachia* strain to inhibit viruses is dependent on the density at which it resides in its host [33-35]. Our previous work demonstrated that *w*Pip resides at a comparable density to *w*Mel in whole *Ae. aegypti* mosquitoes [26]. However, it is not known whether these strains reside differentially within specific tissues that may explain the disparity in their antiviral activity. To test this rigorously we used our Rockefeller *Ae. aegypti* lines carrying the antiviral *Wolbachia* strains *w*Mel or *w*AlbB (classified as *Wolbachia* supergroup A and B strains, respectively), or *w*Pip (from supergroup B), to compare the densities of *Wolbachia* in the salivary glands and midguts of female mosquitoes/line, 6-7 days post emergence. *Wolbachia* density was determined by amplifying the conserved *Wolbachia 16S* rRNA gene and normalising this to the *Ae. aegypti* host *rps17* gene. All 3 *Wolbachia* strains were found to reside at similar densities in whole mosquitoes (between 10 and 14 *Wolbachia* per host cell, Fig. 3A). Given that mosquito salivary glands must become infected with virus in order for the mosquito to transmit DENV, we hypothesized that this tissue would be highly colonized by antiviral *Wolbachia* strains. Surprisingly, the mean relative *w*Mel density was shown to be very low (1 *Wolbachia* per host cell), while *w*AlbB and *w*Pip resided at substantially and significantly higher mean levels (11 and 13 *Wolbachia* per host cell; p<0.0001 Kruskal-Wallis test). Given this unexpected finding, we next examined whether the tissues surrounding the salivary glands may be contributing high *Wolbachia* densities to mediate the antiviral phenotype observed for *w*Mel-carrying mosquitoes. To do this we separated the head/thorax from the abdomen of mosquitoes and determined the *Wolbachia* density. The densities of *w*Mel, *w*AlbB and *w*Pip in the head/thorax closely reflected what was observed in salivary glands alone. It therefore appears that high levels of *Wolbachia* are not required in or around the salivary glands in order to provide an antiviral phenotype. Similarly, the findings for *w*Pip demonstrate that high levels of *Wolbachia* can reside in this critical tissue and not impact virus inhibition. We next used confocal laser scanning microscopy to examine whether *w*Pip localises differently within the salivary gland tissue, to enable DENV replication. Salivary glands were dissected from female mosquitoes 6 days post emergence and stained by fluorescence *in situ* hybridization (FISH) using probes that detect the conserved *Wolbachia* rRNA gene *16S* [11] and DAPI to demarcate the salivary gland tissue. Slides were imaged as 3-dimensional z-stacks and 2D images were generated by Maximum Intensity Projection (MIP) using Fiji software. As expected, there was negligible staining observed in the Rockefeller control samples indicating the specificity of the FISH probes (Fig. 3B). *w*Pip localized similarly to the antiviral strain *w*AlbB, at high levels, but quite diffusely throughout the three lobes of the salivary glands. Consistent with the lower levels of *w*Mel measured by qPCR, this *Wolbachia* strain appeared less prevalent, although interestingly, it was observed to localize in one clustered location of a single lobe. Together, these results show that antiviral *Wolbachia* strains can localise differently within the salivary glands, and that high levels of *Wolbachia* are not required in this tissue to prevent DENV transmission.

**Fig 3.**
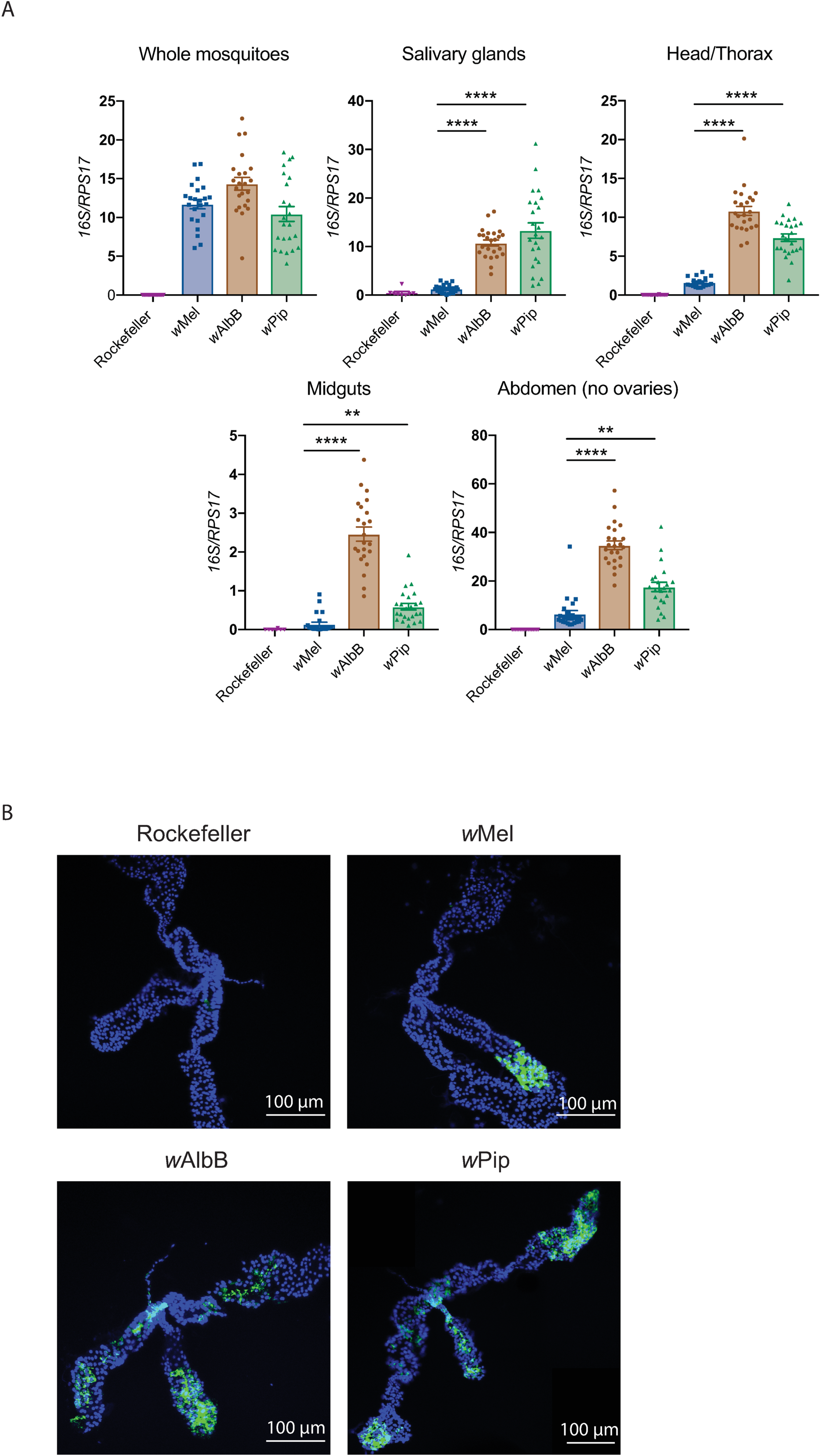
High *Wolbachia* density in *Ae. aegypti* salivary glands and midgut is not required for viral inhibition. (A) Density of *Wolbachia* within female mosquitoes was determined by qPCR following DNA extraction from whole mosquitoes, or from dissected tissues including head/thorax, salivary glands, ovaries, midgut, or abdomen (ovaries removed), as indicated (5-7 days post-emergence). Density is expressed as the mean ratio between the conserved *Wolbachia 16S rRNA* gene and the *Ae. aegypti* host *rps17* gene. Data are the mean and SEM of 24 mosquitoes. Asterisks indicate significance compared to *Ae. aegypti-w*Mel (Kruskal-Wallis test; *p<0.05, ****p<0.0001). B) Salivary glands were dissected from 6 female mosquitoes 6 days post emergence and stained by fluorescence *in situ* hybridization (FISH) using probes that detect the conserved *Wolbachia* rRNA gene *16S* and DAPI to demarcate the salivary gland tissue. Slides were imaged as 3-dimensional z-stacks and 2D images generated by Maximum Intensity Projection (MIP) using Fiji software. Images are representative of >8 salivary gland sets per mosquito line, from 2 independent experiments. Scale bar: 100 µm.

We next examined the density of these *Wolbachia* strains in the midgut of the mosquito (the site of virus adsorption and internalization following an infectious blood meal). Interestingly, *w*Mel was present at very low mean levels in the midgut (0.1 *Wolbachia* per host cell). *w*Pip mean levels were approximately 6 times that of *w*Mel (0.6 *Wolbachia* per host cell), while *w*AlbB resided at the highest density (mean 2.5 *Wolbachia* per host cell; Fig. 3A). To examine whether tissues surrounding the midgut contain high levels of *w*Mel that may explain limited DENV replication in the body of these mosquitoes, we measured the density of each *Wolbachia* strain in mosquito abdomens. Ovaries were removed from the dissected abdomens prior to DNA extraction to prevent obscuring by the high levels of *w*Mel in this tissue [37]. While the *Wolbachia* densities were substantially higher for all lines compared to the midgut alone, the trend across the three lines was almost identical with *w*Mel residing at the lowest density, then *w*Pip, with *w*AlbB residing at the highest density. Thus, the tissues immediately surrounding the midgut are not supporting high levels of *w*Mel to supplement the lower *Wolbachia* densities observed in this tissue.

These findings indicate that a strict localization and density profile at the tissue level is not linked to the antiviral phenotype of *Wolbachia* strains. This conclusion is supported by the recent publication by Flores *et al.* (submitted, PloS Path.) who determined that *w*Mel inhibits DENV replication in mosquito abdomens better than *w*AlbB, despite *w*AlbB residing at higher density than *w*Mel in the midgut and salivary glands as shown here. Of note, *w*AlbB and *w*Pip (both belonging to supergroup B and therefore more closely related to each other than to *w*Mel) seem to display similar tissue distribution and densities in all the tissues examined here, while the profile of *w*Mel is quite different.

### *Wolbachia*-mediated antiviral protection is not dependent on elevated innate immune response pathways in *Ae. aegypti*

We next tested the hypothesis that *Wolbachia* infection in *Ae. aegypti* elevates expression of several innate immune pathway components, particularly in novel *Wolbachia*-host associations, to create an antiviral environment [11, 21-23]. To do this we used our panel of genetically comparable Rockefeller *Ae. aegypti*-*Wolbachia* lines, with the addition of *w*MelPop (also in the Rockefeller background) - a strongly antiviral *Wolbachia* strain that enhances expression of innate immune pathway components in *Ae. aegypti* [11, 21-23]. We assessed the expression levels of Toll pathway components (*cecropin D, cecropin E, defensin C*), known to control anti-DENV defences in mosquitoes [38], and previously reported to be upregulated by some *Wolbachia* strains in *Ae. aegypti* [11, 21-23]. We also assessed expression levels of *C-type lectin* (immune recognition molecule) and *transferrin* (regulation of oxidative stress through iron sequestration), other proteins involved in innate immunity and previously reported to be upregulated by some *Wolbachia* strains in *Ae. aegypti* [11, 21, 23].

Twenty-four female mosquitoes carrying *w*Mel, *w*AlbB, *w*Pip, *w*MelPop, or Rockefeller (no *Wolbachia* control) were collected 6-days post-emergence. Whole mosquito RNA extracts from each individual were converted to cDNA, then expression levels of immune molecules were measured by qPCR, and quantified relative to the mosquito gene *rps17.* Mosquitoes carrying *w*MelPop had significantly elevated levels of each of these representative pathway components compared to the Rockefeller control (Kruskal-Wallis test, Fig. 4). The magnitude of expression increase varied between the transcripts examined, with the smallest increase observed for *C-type lectin* (10-fold increase) and the largest observed for *defensin C* (∼250-fold). In contrast, only very small or no increase in expression of these components was observed in *Ae. aegypti* carrying *w*Mel, *w*AlbB or *w*Pip, relative to the Rockefeller control. *w*Mel did significantly increase expression of *cecropin E* and *defensin C*, but these increases were very small in comparison to *w*MelPop (6- and 2.5-fold increases in the presence of *w*Mel, compared to 180- and 270-fold increases in the presence of *w*MelPop) and were not observed in the other antiviral strain, *w*AlbB. This is the first time *Wolbachia*-induced immune gene expression has been examined in genetically comparable *Ae. aegypti* lines, using a variety of *Wolbachia* strains that either inhibit or do not inhibit DENV. Using this rigorous approach, we can state that elevated expression of these immune components is not consistently associated with a *Wolbachia*-mediated antiviral phenotype in *Ae. aegypti*.

**Fig 4.**
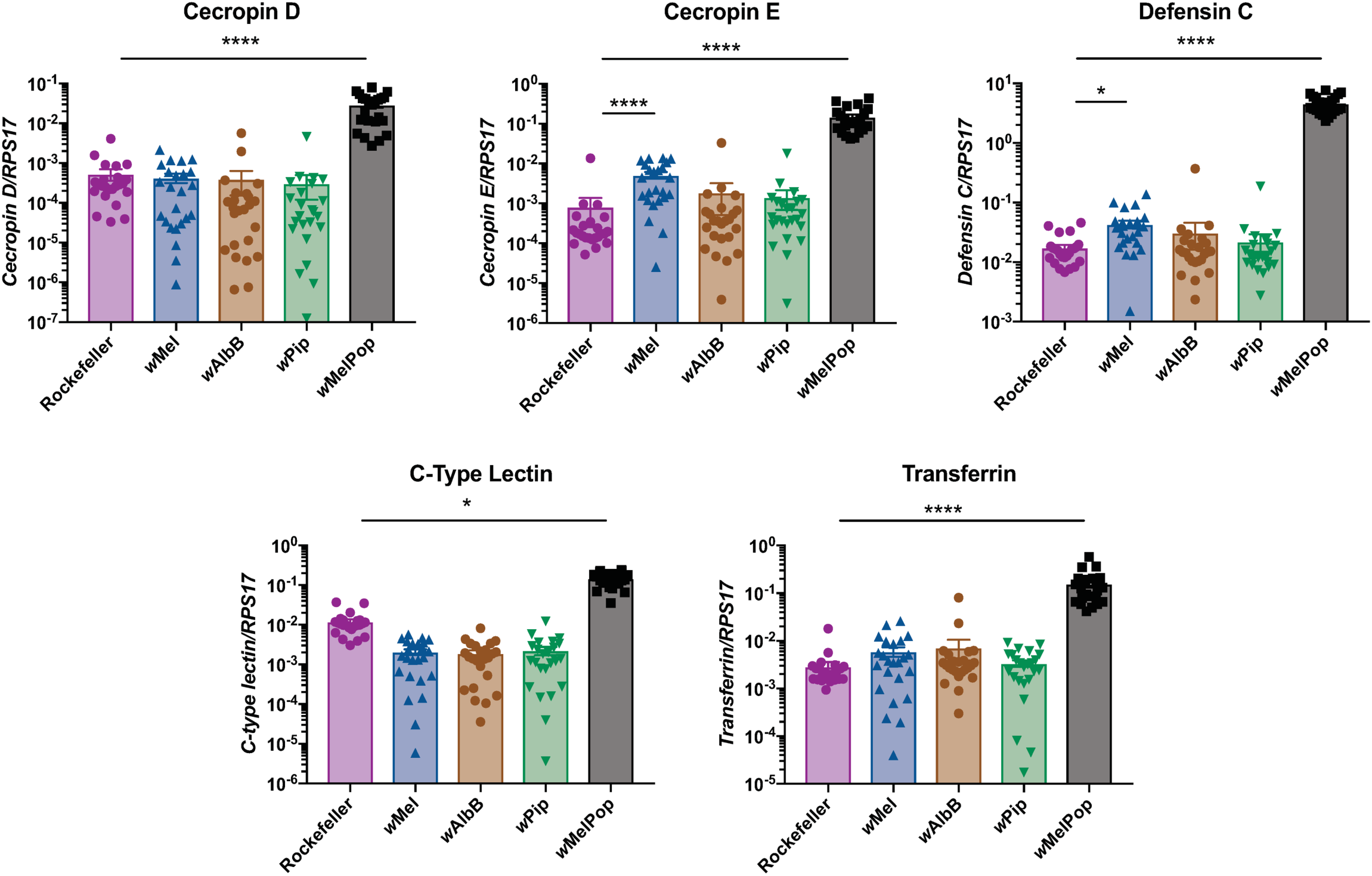
Elevated innate immune response pathways are not required to mediate viral inhibition. Expression levels of selected genes from innate immune pathways were measured by qRT-PCR in 5-day old female mosquitoes, relative to the *Ae. aegypti* host *rps17* gene. Data are the mean and SEM of 24 mosquitoes, representative of 2 independent experiments. Asterisks indicate significance compared to Rockefeller (no *Wolbachia* control) (Kruskal-Wallis test; *p<0.05, ****p<0.0001).

## Discussion

Dissecting the molecular mechanisms that underpin the inhibition of human pathogenic viruses by various *Wolbachia* strains will facilitate the continued success and longevity of *Wolbachia*-based biocontrol programs. This will provide a means to screen for the possible emergence of viral resistance prior to detecting an increase in human disease in *Ae. aegypti*-*Wolbachia* established areas. In addition, it will allow us to identify second-generation *Wolbachia* strains that may restrict viruses using different mechanisms.

*w*Pip is the first *Wolbachia* strain that has been introduced into *Ae. aegypti* without providing antiviral protection towards flaviviruses. A recent study from Ant *et al.*, 2018, introduced *w*AlbA from *Ae. albopictus* into *Ae. aegypti.* This line carried *w*AlbA at a high density yet did not restrict DENV or ZIKV following intrathoracic viral injection [29]. However, when vector competence was examined following an infectious blood meal (DENV or ZIKV), viral infection, dissemination and transmission rates were significantly reduced [30]. We have previously shown that intrathoracic injection challenges with high virus concentrations can overwhelm *Wolbachia*-mediated inhibition [26]. This mode of infection may underestimate the ability of a *Wolbachia* strain to inhibit arboviruses unless injections are performed using a range of virus concentrations. These findings demonstrate the importance of rigorous assay design for vector competence analyses.

Our generation of a novel panel of genetically comparable *Ae. aegypti* lines carrying different *Wolbachia* strains, including one with a non-antiviral strain (*w*Pip), has allowed us to begin rigorously testing a series of hypotheses of the mechanisms underlying viral restriction by *Wolbachia*. Until recently, it was widely accepted that inhibition of viruses in this context correlated with *Wolbachia* density. This dogma was based on a series of experiments using mosquito cell culture and *Drosophila* lines carrying a variety of *Wolbachia* strains, or titrations of a single *Wolbachia* strain by antibiotic treatment [33-35]. However, here, we demonstrate that antiviral phenotype is not dictated by the density at which it resides in either the whole body, salivary glands, the midgut or the tissues immediately surrounding these in the mosquito. This finding builds on, and is consistent with reports that *w*AlbA resides at substantially higher levels than *w*AlbB and *w*Mel in the midgut and the salivary glands of *Ae. aegypti*, despite *w*AlbA showing a relatively limited ability to inhibit flavi- and alphavirus replication [29, 30]. Amuzu and McGraw (2016), examined this in another way, determining that the relationship between DENV inhibition and *w*Mel density within a single *Ae. aegypti* tissue sample is not linear [39]. This finding supports our conclusion that it is not simply the amount of *Wolbachia* present in a tissue that determines whether or not a *Wolbachia* strain is able to inhibit DENV. If *Wolbachia* does not have to be present at high densities to impair viral replication, then this also questions an existing hypothesis that *Wolbachia* impairs viral replication by competing for space within host cells [11, 19, 20]. Further work is needed to analyse the subcellular localisation of various *Wolbachia* strains to determine whether this may determine the antiviral activity of a strain.

Several studies have implied that *Wolbachia* may prime the host innate immune system, preventing arboviral establishment [11, 21-23]. The importance of the contribution of this mechanism has been clouded by the fact that expression levels of these pathway components seems to vary depending on how long the *Wolbachia* strain has resided in its host [21]. We selected a series of immune effectors previously shown to be upregulated by *w*MelPop, *w*Mel or *w*AlbB in *Ae. aegypti*, relative to no-*Wolbachia* control lines. These included toll pathway components (*cecropin D, E*, and *defensin C*), *C-type lectin* (immune recognition molecule), and oxidative stress regulator *transferrin* [11, 21-23]. Using our panel of comparable *Wolbachia*-Rockefeller mosquito lines our findings definitively show that these pathways do not need to be primed at high levels by *Wolbachia* in order to restrict viral replication. While we have not exhaustively considered all innate immune pathways, this finding is consistent with observations from Rances *et al*., 2012, who identified upregulation of many immune genes only occurred in *w*Mel- and *w*MelPop*-Ae. aegypti* where the endosymbiont was a newly acquired infection, but not in the original *D. melanogaster* host where these strains are similarly antiviral [21].

We initially produced the *w*Pip-*Ae. aegypti* line based on work in the native *Culex* mosquito host that showed removal of *w*Pip led to enhanced levels of WNV [32]. Given that *w*Pip does not protect against flaviviruses in *Ae. aegypti*, our results show that there is likely to be a specific *Wolbachia*-host interaction that determines whether a *Wolbachia* strain creates an antiviral state in its host. This host-dependent context has previously been seen for *w*AlbB: this strain does not provide clear protection against flaviviruses in its native *Ae. albopictus* host [13], but effectively inhibits flaviviruses in *Ae. aegypti* [13, 29, 36].

While we are the first to stably introduce *w*Pip into *Ae. aegypti, w*Pip was previously introduced into *Ae. albopictus:* as a single infection, as a double infection with *w*Mel [40], and as a triple infection with the natural *w*AlbA*w*AlbB *Wolbachia* strains [41]. These studies have shown that *w*Pip alone in *Ae. albopictus* is not antiviral, while combining *w*Pip with *w*Mel does inhibit arboviruses, and the addition of *w*Pip to the natural *w*AlbA/*w*AlbB combination significantly reduces DENV and ZIKV replication. This complicated scenario may suggest that these *Wolbachia* strains each modify particular aspects of the host which together create an antiviral state [42].

This finding poses an interesting issue for selecting *Wolbachia* strains for novel *Ae. aegypti* lines – that is, how can we predict whether a new strain will induce an antiviral state? Notably, so far it seems that the magnitude of the antiviral effect of a *Wolbachia* strain measured in *Drosophila spp*. predicts how the strain will behave in *Ae. aegypti* (from the 5 strains examined) [10, 11, 26, 28, 29, 43, 44]. This has not been the case for mosquito-derived *Wolbachia* strains – *w*AlbA and *w*AlbB are not antiviral in their native host, but do provide protection in *Ae. aegypti*, while *w*Pip is reported to be antiviral in *Cu. quinquifaciatus*, but not in *Ae. aegypti* [13, 30, 32]. Perhaps this is due to differences in the way these *Wolbachia* strains localise in each host. And/or, if multiple mechanisms contribute to viral inhibition, perhaps each host-*Wolbachia* combination has some or all of these mechanisms in play.

The introduction of other novel *Wolbachia* strains into *Ae. aegypti* is required to determine if this trend holds.

In this study we have generated a unique and powerful tool: a panel of *Ae. aegypti* lines that differ only in the *Wolbachia* strain that they carry, including *w*Pip-*Ae. aegypti* which does not restrict flavivirus replication, dissemination or transmission. This is an important finding as it identifies a refined negative control for studies trying to understand how *Wolbachia* affects its host. A *Wolbachia*-free control has been used in the past, but this control does not account for the many host effects *Wolbachia* can induce just by residing as an endosymbiont, that may not be responsible for creating an antiviral state [45, 46]. The strength of this approach was recently shown by LePage *et al*., who compared the genomes of *Wolbachia* strains that do or do not induce CI to manipulate host reproduction, identifying the two genes responsible for inducing this phenotype [47]. We have now initiated a series to experiments to carefully dissect the mechanisms that drive *Wolbachia*-mediated viral inhibition.

## Methods

### Mosquito rearing

All *Ae. aegypti* mosquitoes were reared and maintained as described previously [26, 27, 37]. Briefly, adult mosquitoes were maintained at 26°C, 65% relative humidity (RH) and a 12 h light:dark cycle in a climate-controlled room. Mosquitoes were blood fed on the arms of human volunteers (Monash University human ethics permit CF11/0766-2011000387). The *Wolbachia*-infected *w*Mel, *w*AlbB and *w*Pip lines as well as matched Tet-control lines (*Wolbachia* infected lines that have been cured of their infection by tetracycline treatment) used in these experiments have been described previously [26, 28, 48] [Flores *et al.*, submitted PLOS Path].

To generate a panel of genetically comparable *Wolbachia*-carrying *Ae. aegypti* lines, we backcrossed females from the *w*Mel, *w*AlbB, *w*MelPop and *w*Pip to males of the inbred laboratory *Ae. aegypti* line, Rockefeller [49] (BEI resources), for six generations.

To exclude any influence of mosquito age on our experiments, age-controlled adults emerging within a 24 h window were used.

### Vector competence

DENV-3 Cairns 08/09 strain (Genbank accession number: JN406515.1) and DENV-2 strain (originally isolated from a patient in Vietnam in 2010) were prepared by inoculation of C6/36 cells with a multiplicity of infection (MOI) of 0.1 and collection of culture supernatant 6-7 days later. KUNV stocks (Genbank accession number: MRM61C) were prepared by inoculation of C6/36 cells with a multiplicity of infection (MOI) of 1 and collection of culture supernatant 48 hours later. Virus concentrations were determined by TCID_50_ as previously described [50] using monoclonal antibody 4G2 [51].

For feeding experiments with DENV-3 (Cairns 08/09) infected blood, 100 seven-day old age-controlled female mosquitoes were placed in 500 mL plastic containers (five containers per *Wolbachia* line, three containers per Tet line), starved for up to 24 h and allowed to feed on a 50:50 mixture of defibrinated sheep blood and tissue culture supernatant containing freshly harvested 6.6 x 10^6^ TCID_50_/mL of DENV-3. Feeding was done through a piece of desalted porcine intestine stretched over a water-jacketed membrane feeding apparatus preheated to 37°C. Mosquitoes were left to feed in the dark for approximately 1-2 hours. Fully engorged mosquitoes were placed in 500 mL containers at a density of < 25/container, and incubated for 15 d at 26°C with 65% RH and a 12 h light/dark cycle.

For adult microinjections, 60 six- or seven-day old age-controlled female mosquitoes were anesthetized by CO_2_. Mosquitoes were injected intrathoracically with 69 nL of DENV (DENV-3 Cairns 08/09 strain at 6.3 ×10^5^ or 6.3 ×10^4^ TCID_50_/ml, or DENV-2 Vietnam strain at 2.4 × 10^5^) in RPMI media (Life Technologies) using a pulled-glass capillary and a handheld microinjector (Nanoject II, Drummond Scientific). For KUNV injections, 69 nL of 1.4 ×10^7^ or 1.4 ×10^6^ TCID_50_/ml was injected per mosquito using the same method as described for DENV-3. Injected mosquitoes were incubated for 7 days (15 mosquitoes/cup) at 26°C with 65% RH and a 12 h light/dark cycle.

To quantify DENV-3 or KUNV genomic copies, total RNA was isolated from mosquitoes (entire mosquitoes for injection experiments, or head and bodies separately for blood-fed mosquitoes) using the RNeasy 96 QIAcube HT kit (Qiagen). DENV-3 RNA was amplified by qRT-PCR (LightCycler Multiplex RNA Virus Master, Roche), using primers to the conserved 3’UTR: Forward 5’-AAGGACTAGAGGTTAGAGGAGACCC; Reverse 5’-CGTTCTGTGCCTGGAATGATG; Probe 5’-HEX-AACAGCATATTGACGCTGGGAGAGACCAGA-BHQ1-3’ [52]; absolute copies were determined by extrapolation from an internal standard curve generated from plasmid DNA encoding the conserved 3’UTR sequence. Mosquito extracts with ≥1000 copies of DENV per body were scored positive, based on the LOD_95_ (limit of detection 95%) for DENV-3 with this primer set. KUNV RNA was amplified by using primers that span the 3’ end of the conserved NS5 gene, and the 3’UTR: Forward 5’-AACCCCAGTGGAGAAGTGGA; Reverse 5’-TCAGGCTGCCACACCAAA; Probe 5’-HEX - CGATGTTCCATACTCTGGCAAACG -BHQ1-3’ [53]. KUNV RNA copies were quantified relative to *Ae. aegypti* house-keeping gene *rps17* using the delta CT method (2^CT^(reference)/ 2^CT^(target)). KUNV RNA copies with a CT of < 33 were scored positive for infection. Note that all surviving mosquitoes were processed for virus injection experiments, while a maximum of 72 mosquitoes/line were collected and processed for blood feeding experiments.

### Virus transmission

To determine whether *Wolbachia*-infected mosquitoes were capable of transmitting infectious virus, 15 blood-fed mosquitoes per line were collected at 15 days post blood feed (donor mosquitoes). The proboscis from each donor was inserted into a 10 μL pipette tip containing 10 μL 1:1 FBS:30% sucrose [54]. Legs and wings were removed to encourage the mosquitoes to spit. Pipette tips were collected 1 hour later and the virus solution ejected onto parafilm. The solution was drawn up into a pulled-glass capillary attached to a handheld microinjector (Nanoject III, Drummond Scientific) and 600 nL was injected into 6 seven-day old recipient *w*Mel.Tet mosquitoes. Replicate recipient mosquitoes were stored in a single container for 7-days post injection. RNA was extracted from the whole bodies of all surviving mosquitoes, and qRT-PCR was performed as described above.

### Wolbachia density and distribution

Relative *Wolbachia* density in *w*Mel, *w*AlbB and *w*Pip was determined in whole or dissected tissues from female mosquitoes at 5-days post emergence, using qPCR with primers to amplify a fragment of the conserved *16S rRNA* gene (forward primer: 5’-GAGTGAAGAAGGCCTTTGGG-3’, reverse primer: 5’-CACGGAGTTAGCCAGGACTTC-3’, probe 5’ LC640-CTGTGAGTACCGTCATTATCTTCCTCACT-IowaBlackRQ-3’) and the reference *Ae. aegypti rps17* gene (forward primer: 5’-TCCGTGGT ATCTCCATCAAGCT-3’, reverse primer: 5’-CACTTCCGGCACGTAGTTGTC-3’, probe 5’FAM CAGGAGGAGGAACGTGAGCGCAG-BHQ1-3’) [26]. *Wolbachia* densities were quantified relative to *rps17* using the delta CT method as previously (2^CT^(reference)/ 2^CT^(target)).

### Salivary gland dissections and fluorescence in-situ hybridization (FISH) staining

Female mosquitoes were collected 6-days post emergence, knocked down at -20°C for 2 minutes, then kept in a petri dish on ice until dissection. Individuals were dissected on a microscope slide in a drop of PBS. Briefly, the head of the mosquito was sliced off using a dissection needle, and the salivary glands popped out by gently squeezing the thorax with needle-tipped forceps. Salivary glands were gently transferred to a small droplet of PBS on poly-lysine-coated slides. Tissues were fixed in cold 4% paraformaldehyde in PBS for 15 minutes, rinsed 3 times in PBS, then permeabilised in 100% ethanol for 5 minutes and air dried. Slides were incubated in hybridization buffer containing fluorescently labelled *16S rRNA* probes (cross-reactive with all three *Wolbachia* strains) [11] overnight at 37°C in a humidified chamber. Slides were washed in SSC buffers + 10 mM DTT, stained with DAPI, and mounted as described by Moreira *et al.*, 2009 [11].

### Confocal microscopy

Slides were imaged using a Nikon C1 Upright confocal microscope at 20X magnification (under oil) as 3-dimensional z-stacks with a step-size of 3 microns. Images were acquired with NIS-Elements software. Maximum Intensity Projection images and scale bars were generated in Fiji software (Version 1.52; National Institutes of Health). Note that the image of *w*Pip salivary glands was produced by stitching together two images taken from the same sample, as the lobes spread too wide to image under a single field of view [55].

### Quantitative RT-PCR for immune gene targets

RNA was extracted from 6-day old female mosquitoes (24 per line including Rockefeller, *w*Mel, *w*AlbB, *w*Pip) using the RNeasy 96 QIAcube HT kit (Qiagen). Samples were DNAseI treated and cDNA was generated from 8 μL of purified RNA/individual (∼500 ng of RNA) using SuperScript™ III First-Strand Synthesis System (ThermoFisher). cDNA was diluted 2-fold with RNase-free water then expression levels of selected immune genes was determined by amplifying 1 μL of cDNA with primers for *cecropin D* (AAEL000598; forward 5’-GCTAGGTCAAACCGAAGCAG, reverse 5’-TCCTACAACAACCGGGAGAG) [23], *cecropin E* (AAEL000611; forward 5’-TTGCACTCGTTCTGCTCATC, reverse 5’-ACACGTTTTCCGACTCCTTC) [23], *defensin C* (AAEL003832-RA; forward 5’-GCTGAGTGGGTTCGGTGTAG, reverse 5’-CGCGTTACAATAGCCTCCTC) [21], *C-type lectin* (AAEL005641; forward 5’-GTCTCCGGGTGCAATACACT, reverse 5’-CCCTATCGTTCCACTTCCAA) [23] or *Transferrin* (AAEL0015458; forward 5’-TCAGGATCTGATGGCCAAAC, reverse 5’-GCCTTGACCTTCTCCAGACA) [23]. Expression levels were normalized to the *Ae. aegypti* house-keeping gene *RPS17* (forward 5’-TCCGTGGT ATCTCCATCAAGCT, reverse 5’-CACTTCCGGCACGTAGTTGTC) using the delta CT method (2^CT^ (reference)/ 2^CT^ (target)).

## Ethics statement

Blood feeding by volunteers (Monash University human ethics permit no CF11/0766-2011000387) for this study was approved by the Monash University Human Research Ethics Committee (MUHREC). All adult volunteers provided informed written consent; no child participants were involved in the study.

## Figure Legends

**Fig S1.**
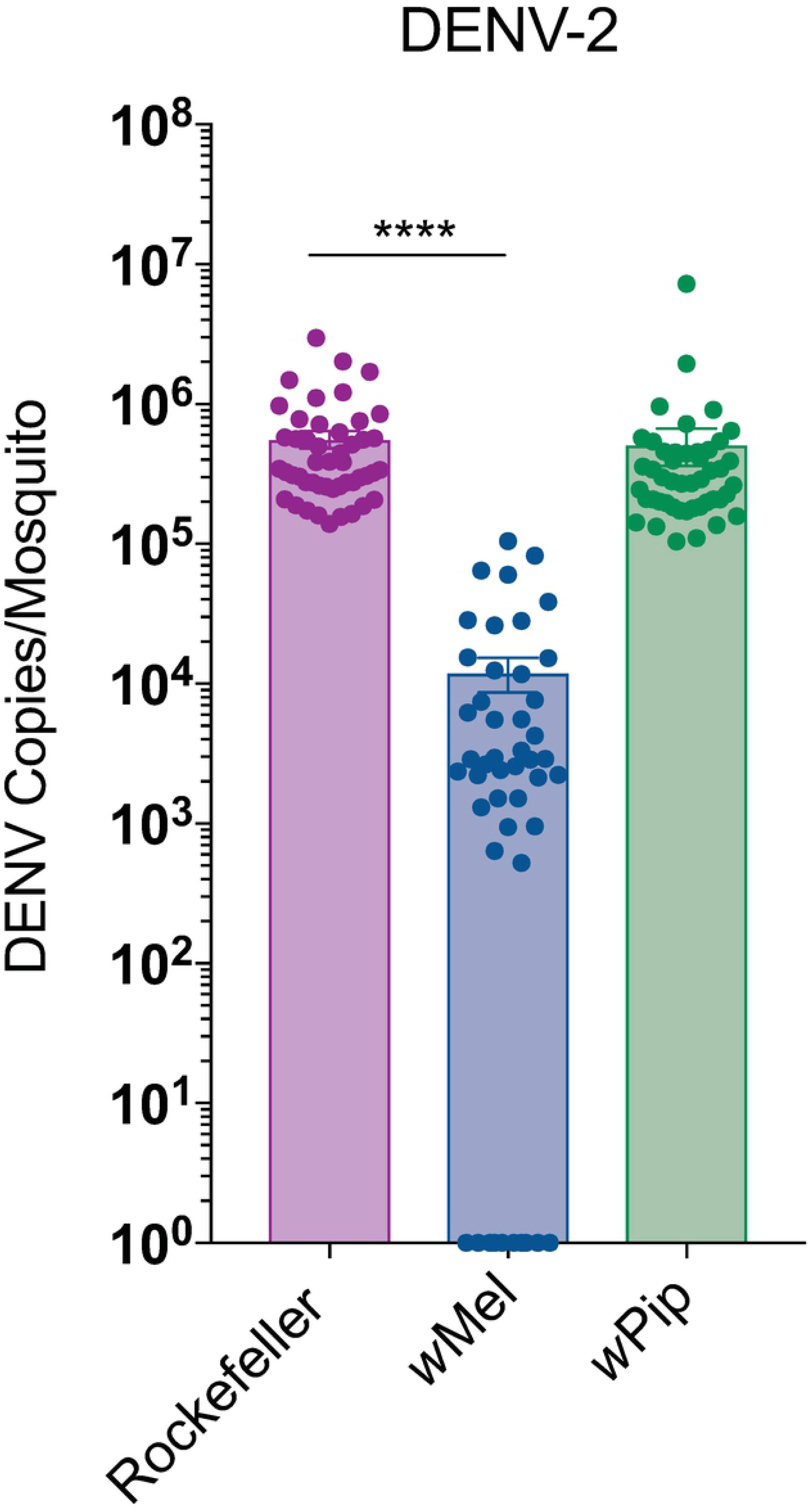
DENV-2 (isolated from a patient in Vietnam in 2010) was injected into the thorax of 7-day old female mosquitoes at 2.4 ×10^5^ TCID_50_/ml. RNA was extracted from whole mosquito bodies 7-days post infection and virus replication was quantified by qRT-PCR. Data are the mean number of DENV genome copies per mosquito ± SEM with individual data points overlaid. Asterisks indicate significance compared to Rockefeller (no *Wolbachia* control) (Kruskal-Wallis test; ****p<0.0001).

## Acknowledgements

We wish to thank Etiene C. Pacidônio, Daniela S. Gonçalves, Kimberley Dainty, Ritzel Gimeno and Elvina Lee for technical assistance. We thank Professor Roy Hall (University of Queensland) for provision of the 4G2 antibody, and Professor Jason Mackenzie (University of Melbourne) for gifting the Kunjin virus stock. The authors acknowledge Monash Micro Imaging, Monash University, for the provision of instrumentation, training and technical support. The following reagent was obtained through BEI Resources, NIAID, NIH: *Aedes aegypti*, Strain ROCK, MRA-734, contributed by David W. Severson. This work was supported by a Wellcome Trust Grant (212914/Z/18/B), and National Health and Medical Research Council Australia Ideas Grant APP1182432.

